# Gait variability is affected more by peripheral artery disease than by vascular occlusion

**DOI:** 10.1101/2020.10.21.348433

**Authors:** Hafizur Rahman, Iraklis I. Pipinos, Jason M. Johanning, Sara A. Myers

**Affiliations:** Department of Biomechanics, University of Nebraska at Omaha, Omaha, Nebraska, USA; Department of Surgery, Veterans’ Affairs Medical Center of Nebraska and Western Iowa, Omaha, Nebraska, USA; Department of Surgery, University of Nebraska Medical Center, Omaha, Nebraska, USA

**Author notes:** Corresponding author, (SAM).

## Abstract

**Background:** Patients with peripheral artery disease (PAD) have altered gait variability from the first step they take, well before the onset of claudication pain. The mechanisms underlying these gait alterations are poorly understood.

**Aims:** This study sought to isolate the effect of reduced blood flow on gait variability by comparing healthy older controls and patients with PAD. We also determined the diagnostic value of gait variability parameters to identify the presence of PAD.

**Methods:** Thirty healthy older controls and thirty patients with PAD walked on a treadmill at their self-selected speed in rested (normal walking for controls; pain free for PAD) and reduced blood flow (post vascular occlusion with thigh tourniquet for controls; pain for PAD) conditions. Gait variability was assessed using the largest Lyapunov exponent, approximate entropy, standard deviation, and coefficient of variation of ankle, knee, and hip joints range of motion. Receiver operating characteristics curve analyses of the rested condition were performed to determine the optimal cut-off values for separating individuals with PAD from those without PAD.

**Results and Discussion:** Patients with PAD have increased amount of variability for knee and hip ranges of motion compared with healthy older group. Comparing for conditions, reduced blood flow demonstrated increased amount of variability compared with rested blood flow. Significant interactions between group and condition occurred at the ankle for Lyapunov exponent, approximate entropy, and coefficient of variation. A combination of gait variability parameters correctly identifies PAD disease 70% of the time or more.

**Conclusions:** Gait variability is affected both by PAD and by the mechanical induction of reduced blood flow.

## 1. Introduction

Peripheral artery disease (PAD) is a common cardiovascular disorder that is associated with poor health outcomes, immobility, and physical dependence [1–3]. The most common symptom is intermittent claudication, a cramping pain that occurs in the calves, thighs and/or buttocks brought on by physical activity and relieved with rest. Patients with PAD are also significantly compromised by decreased physical activity and a higher prevalence of falls [4–6]. Advanced biomechanical evaluations have been undertaken to elucidate the mechanisms responsible for these problems [7–12]. The results of these works demonstrate that the gait changes are driven by pathophysiological alterations that occur long before intermittent claudication symptoms drive patients to see a physician for pain during walking [6,13,14].

With twenty-nine percent of individuals over the age of 70 years being affected by PAD, the older population is particularly at risk for developing PAD [15]. Older individuals experience in less severe levels similar gait problems as patients with PAD, including decreased strength, decreased push-off kinetics, and higher rates of falling [4,5,9,16]. Risk factors for falling in older individuals are similar as those for patients with PAD, including gait and balance impairments [17]. The incidence of PAD in older individuals makes it important to dissect the different pathophysiologic mechanisms that contribute to the falls and mobility problems in elders with PAD compared with the mechanisms operating in the normal aging process. Further, a minimally invasive method that allows the detailed measurement of the severity of functional compromise in PAD and the monitoring of changes in the disease progression would also be a useful clinical tool.

Gait variability refers to the natural alterations that occur within and across gait cycles. The measurement of gait variability is a method that has shown promise in predicting falls, distinguishing between pathological and healthy individuals, and as an indicator of overall health of the biological system [18,19]. Traditionally, movement variability was thought of as “noise”, but the current consensus is that it is inherent within all biological systems [20,21]. The idea of “healthy variability” was originally put into application in the study of heart rhythm, with healthy heart rates exhibiting an ordered but varying pattern described using mathematical scaling techniques or fractal analysis [22]. It is thought that the ordered variations in the steady state output of healthy systems represent the underlying physiologic capability to perform flexible adaptations to everyday and second by second stresses placed on the human body [23–25]. Consequently, the normal variability of human gait is the product of continual adjustments to the always changing environmental and morphological conditions affecting normal walking.

Alterations in gait variability have been found in numerous pathological populations, with some exhibiting deviations towards less order and others more order in gait [26–32]. Previous studies have identified that patients with PAD have a significant increase in gait variability as compared with healthy controls, indicating a less ordered pattern and significant deterioration of the locomotor system [7,8]. The specific mechanisms leading to gait variability alterations from the “optimal” levels have not been elucidated. The muscle discomfort of intermittent claudication is commonly thought to be the factor that is responsible for the mobility problems in patients with PAD [33], but in both gait biomechanics and gait variability studies, the patients have significantly altered gait from the first step they take and well before pain onset [7,8]. Previous work from our and other groups points to abnormal lower extremity blood flow and underlying cellular abnormalities in the lower extremity muscles and nerves as possible mechanisms contributing to differences in the gait biomechanics and variability patterns as compared to healthy individuals [34–43].

Previous work from our group determined the role of walking-induced ischemia and its related muscle pain symptoms in the alterations of gait variability in patients with PAD by examining healthy younger and healthy older individuals before and after an induced vascular occlusion using a tourniquet placed at the level of the subject’s thigh [44,45]. Both studies indicated gait variability was significantly altered during the post-vascular occlusion condition. However, the magnitude of changes post occlusion were less than the magnitude of gait variability alterations seen in patients with PAD. The average increases in gait variability from baseline to post occlusion for the healthy younger and older groups were 21% and 11.9%, respectively [44,45]. In comparison, patients with PAD have gait variability values that are 48% higher than healthy younger and 34% higher than healthy older individuals [8].

The current study builds on our previous work by directly comparing gait variability between healthy older controls and patients with PAD during rested and reduced blood flow conditions. The rested conditions consisted of normal walking in the healthy group, and the pain free condition in the PAD group, during which the patient walks prior to the onset of ischemic claudication pain. The reduced blood flow conditions included the post occlusion condition in the healthy group, and the pain induced condition in the PAD group, during which the patient walks prior to the onset of ischemic claudication pain. We hypothesized that gait variability values would be higher in patients with PAD compared to controls during both conditions. We also hypothesized that gait variability values would be increased during the reduced blood flow condition as compared to the rested condition for both control and patients with PAD.

The current screening method for PAD is through measurement of ankle-brachial index in a vascular laboratory. This test is considered a specialized test that requires a separate medical appointment. Gait variability has potential to be assessed remotely in a clinical or home setting. It may provide an early indicator of functional problems, assess improvements following treatments, or provide an alert for declines that can occur with disease progression. To test the measurement of gait variability as a potential clinical tool, we also established threshold gait variability values which indicate the presence of PAD using a receiver operating characteristics curve. We hypothesized gait variability would provide acceptable discrimination between individuals with PAD and those without PAD.

## 2. Materials and methods

### 2.1 Participants

Thirty healthy older participants (age: 60.1 ± 8.03 years, mass: 86.6 ± 16.1 kg, height: 176.7 ± 8.3 cm, gender: 25 males, 5 females) and thirty symptomatic patients with PAD with moderate arterial occlusive disease and bilateral claudication (age: 63.8 ± 9.10 years, mass: 81.1 ± 14.7 kg, height: 171.8 ± 5.1 cm, gender: 28 males, 2 females) participated in the study. Prior to data collection informed consent was obtained from all participants according to the guidelines of our Institutional Review Boards.

Patients were screened and evaluated by one of two board-certified vascular surgeons. The evaluation included a detailed history and physical examination. Patients with PAD were excluded if they experienced pain or appreciable limitation during walking for any reason other than claudication. Such significant limitations included cardiac, pulmonary, neuromuscular, or musculoskeletal or joint disease. Control subjects had an ankle-brachial index ≥ 1.0 and no subjective or objective ambulatory dysfunction. Controls were excluded if they experienced any discomfort or limitation during walking. All subjects had their gait analyzed in the biomechanics laboratory.

### 2.2 Experimental procedures and data collection

Following informed consent, height, body mass, and anthropometric measures were taken for all subjects. Healthy older individuals had lower extremity blood flow measured by taking the systolic pressures at the brachial artery in the arm and the dorsal pedis and posterior tibial arteries at the ankle to confirm acceptable ankle-brachial index values. Patients with PAD had ankle-brachial index values assessed in the same manner, however these measures were taken at the respective clinical site and did not need to be repeated in the biomechanics laboratory. Reflective markers were placed on specific anatomical locations as previously described [8]. Participants were then allowed to get familiar with walking on the treadmill (BodyGuard Fitness, St. Georges, QC, Canada). During familiarization, subjects were asked to select a comfortable walking speed, which was then identified as the self-selected walking speed and was used for all further testing.

A Motion Analysis eight camera (Eagle cameras, Motion Analysis Corp, Santa Rosa, CA) system was used to capture kinematics while participants walked on the treadmill. Three-dimensional movements were acquired at 60 Hz using EVART software (Motion Analysis Corp, Santa Rosa, CA). Prior to recording the kinematics data, subjects stood in a calibration device to collect an anatomical reference position of each leg. This position was used to orient the lower extremity segments and served as the reference point for relevant joint angles. Data collection continued as follows for the healthy older controls and for patients with PAD:

#### 2.2.1 Healthy older controls

Participants walked on the treadmill at their self-selected pace for three minutes, which was considered as the baseline condition. Next, vascular occlusion was induced by placing thigh cuffs (Omron® Exactus Aneroid Sphygmomanometer Model 108MLNL, Kyoto, Japan) bilaterally on the upper thighs and occlusion tourniquets (CyberTech™ Mechanical Advantage Tourniquet MAT01, Maharashtra, India) just above the knee while subjects stood on the treadmill. The cuffs were inflated to 200 mm Hg and maintained for three minutes. The chosen level of pressure and time of occlusion is standard and has been used in previously published similar studies to induce ischemia in the legs [46–51]. After three minutes of occlusion, the thigh cuffs were removed, and the subjects immediately began walking on the treadmill. Three minutes of treadmill walking was recorded for post vascular occlusion and defined as the post condition, although only the first 3300 data points were used for analysis.

#### 2.2.2 Patients with PAD

Participants walked on a treadmill at self-selected speed for three minutes or until the onset of claudication pain, whichever came first. This was the pain free condition. Subjects were then required to rest for a minimum of 10 minutes to ensure that they were relieved from claudication pain. Next, the treadmill was inclined to 10% grade and the subjects walked until the onset of claudication pain. Once pain was present, the treadmill was lowered to 0% grade and the patients walked on the level treadmill with claudication pain for as long as tolerable up to three minutes. This was defined as the pain condition.

### 2.3 Data analysis

Coordinate trajectory data of each marker were exported and processed using custom codes in MATLAB software (MathWorks Inc, Natick, Mass). This software was used to calculate relative joint angle time series from the kinematic data for the ankle, knee, and hip for all trials as previously described [10,45]. Joint kinematic variability has been shown to be a more sensitive measure of differences between groups than variability of stride characteristics (stride time, step time) [52]. All trials were cropped to 3300 data points, which is long enough to allow 30 continuous footfalls and is considered adequate for nonlinear analysis [53,54]. We assessed gait variability using both linear and nonlinear analysis.

#### 2.3.1 Gait variability: linear analysis

The linear analysis gives information regarding the amount of variability present in gait patterns and is used in conjunction with the nonlinear analysis [54]. Ranges of motion of the ankle, knee, and hip joint angles were calculated for the gait cycles in every trial. Means, standard deviations, and coefficients of variation ((standard deviation / mean) * 100) were then calculated for each variable. These calculations were made using custom written laboratory codes in MATLAB software (MathWorks Inc, Natick, MA, USA).

#### 2.3.2 Gait variability: nonlinear analysis

The largest Lyapunov exponent and approximate entropy were utilized for this study. Linear analysis considers a few specific points in the series, but both the largest Lyapunov exponent and approximate entropy investigate the temporal structure of the entire time series of the joint angle. The largest Lyapunov exponent approximates the sensitivity of the locomotor system to perturbations by quantifying the exponential separation of nearby trajectories in the reconstructed state space of the joint angle time series. As nearby points of the state space separate, they diverge rapidly and can produce instability. The largest Lyapunov exponent ranges from zero in a stable system with little to no divergence (e.g., sine wave) to a larger value around 0.5 for an unstable system with a high amount of divergence (e.g., white noise) [54–56]. A deterministic or ordered signal that demonstrates mathematical chaos will have a largest Lyapunov exponent value between zero and 0.5 when it is calculated using the Wolf et al algorithm implemented in the Chaos Data Analyzer [57]. To numerically calculate the largest Lyapunov exponent for each joint angle time series for each trial the *Chaos Data Analyzer* professional version (American Institute of Physics, Raleigh, NC, USA) was used [58]. The detailed description of actual calculation of the largest Lyapunov exponent was previously described [8,44,45]. No surrogation was performed in this experiment, as it has been performed for these groups previously [7,8,44,45].

Approximate entropy was calculated to determine the repeatability present in each trial [26,54,59]. Approximate entropy gives information regarding the regularity of a time series by measuring the logarithmic probability that a series of data points at a certain distance apart will exhibit similar relative characteristics on the next incremental comparison with the state space [60–63]. Time series with a greater likelihood of remaining the same distance apart in the next incremental comparison with a state space (periodic and predictable) will result in approximate entropy values close to zero. Those data points that vary widely in distances between data points (irregular and not predictable) will result in high values close to two. For example, the approximate entropy value for a periodic time series like the sine wave will be close to zero and the value for a random time series like white noise will be close to two. A signal that demonstrates mathematical chaos will lie somewhere between zero and two. A more detailed description of the calculation of the approximate entropy was previously described [8,44,45].

#### 2.3.3 Receiver operating characteristics (ROC) curve analysis

Mean values for linear and nonlinear gait variability parameters from the pain free condition for patients with PAD and from the baseline condition for healthy older controls were utilized to construct receiver operating characteristics curves. First, logistic regression was performed independently for each gait variability dependent variable to determine if each variable was significantly associated with disease status (presence or absence of PAD). Next, the ROC curves were calculated for all variables with a significant relationship with disease status. The area under the curve, which gives the variable’s overall ability to predict the presence of PAD was calculated. Area under the curve values range from 0.0, which represents no ability to discriminate PAD from healthy individuals, to 1.0, which would give perfect discrimination of those with PAD from those without PAD [64–66]. The level of acceptable discrimination was set at 0.75 (75% of the area under the curve) for each variable [64,66]. Prediction equations were calculated for all dependent variables that had area under the curve values greater than or equal to 0.75. Optimal cut-off points, percentage correct, sensitivity, and specificity values were calculated to provide information regarding the accuracy of each prediction equation. The probability of the presence of PAD for an individual observation (π(x)) is calculated from each prediction equation using the form of Equation (1), which is derived from the *logit* transformation. If π(x) is greater than the optimal cut-off point for the dependent variable, then the observation is classified as PAD. If π(x) is less than the cut-off point, the observation is classified as healthy.

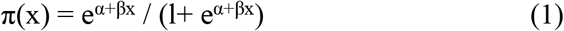

### 2.4 Statistical analysis

Group and condition means for the standard deviation and the coefficient of variation of the range of motion, largest Lyapunov exponent values, and approximate entropy values were calculated for the ankle, knee, and hip joint angles. For each dependent variable, a two by two mixed ANOVA was used to detect differences for group (PAD versus healthy older control) and condition (baseline/pain free versus post vascular occlusion/pain) factors. When a significant interaction was identified, Tukey tests were used for post-hoc analysis to identify significant differences between the group/condition combinations [67]. Statistical comparisons were performed using SPSS software (version 26, IBM, Armonk, NY). The level of significance was set at 0.05.

## 3. Results

### 3.1 Linear measures of variability

There were several differences between groups (healthy older control versus patients with PAD). Patients with PAD exhibited increases in standard deviation and coefficient of variation values of the knee and hip ranges of motion as compared to healthy older controls (*p* < 0.05; Figs 1 and 2). There were also significant differences for condition (baseline/pain free versus post occlusion/pain). Specifically, the post occlusion/pain condition had increases in standard deviation values of the knee and hip ranges of motion compared to the baseline/pain free condition.

**Fig 1.**
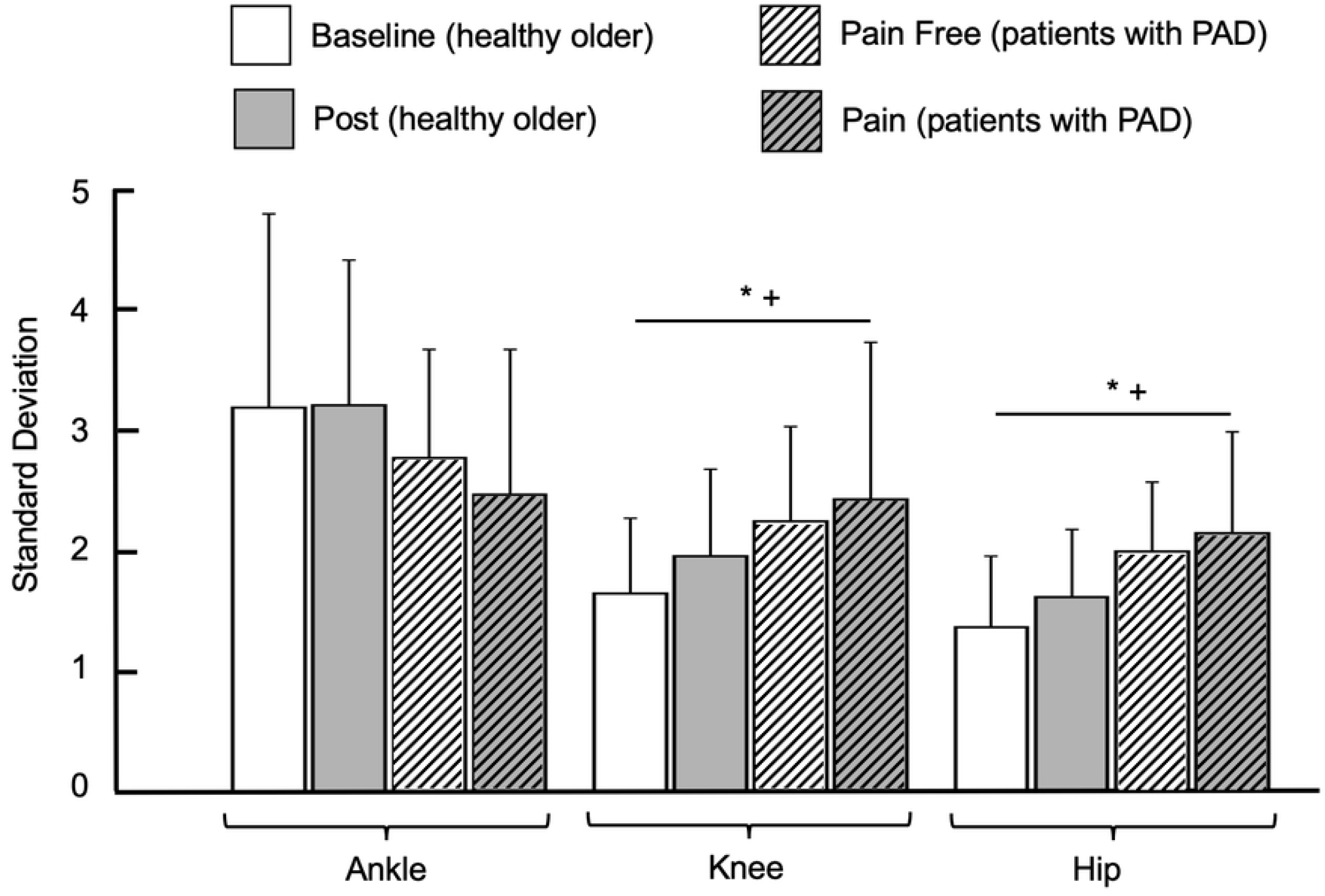
Standard deviation for healthy older individuals in the baseline and post vascular occlusion (post) conditions and for patients with peripheral artery disease (PAD) in the pain free and pain conditions for ankle, knee, and hip ranges of motion. * indicates significant differences between conditions (Baseline/Pain Free vs. Post/Pain, *p* < 0.05). + indicates significant differences between groups (PAD vs. healthy older,*p* < 0.05). # indicates post-hoc tests while significant interaction exists between groups and conditions. Bar graphs represent the mean values and error bars represent the standard deviation.

**Fig 2.**
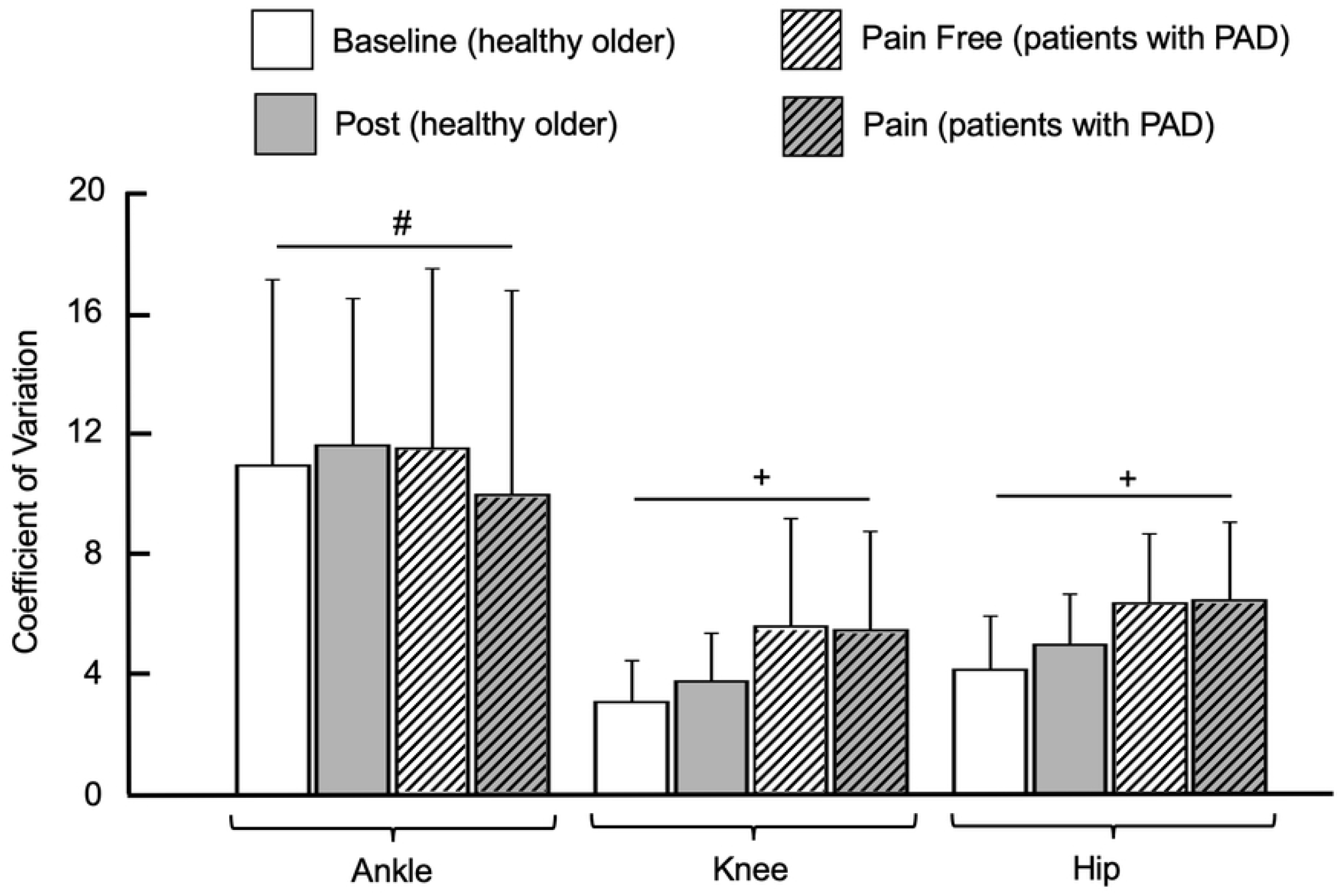
Coefficient of variation for healthy older individuals in the baseline and post vascular occlusion (post) conditions and for patients with peripheral artery disease (PAD) in the pain free and pain conditions for ankle, knee, and hip ranges of motion. * indicates significant differences between conditions (Baseline/Pain Free vs. Post/Pain, *p* < 0.05). + indicates significant differences between groups (PAD vs. healthy older,*p* < 0.05). # indicates post-hoc tests while significant interaction exists between groups and conditions. Bar graphs represent the mean values and error bars represent the standard deviation.

There was also a significant interaction between group and condition for the ankle range of motion coefficient of variation; patients with PAD exhibiting increased values in the pain free condition as compared with the pain condition (Fig 2). There were no other significant interactions between group and condition means. Collectively, the results of the linear analysis indicate that the PAD group exhibited increased amount of variability for the knee and hip ranges of motion compared with the healthy older group. Additionally, the reduced blood flow condition (post occlusion in the healthy older and pain in the PAD) demonstrated increased amount of variability for the knee and hip ranges of motion compared with the rested blood flow condition (baseline/pain free).

### 3.2 Nonlinear measures of variability

Regarding the effect of group, patients with PAD showed increased values of the largest Lyapunov exponent for the ankle, knee, and the hip joint time series (*p* < 0.05; Fig 3), but there were no significant group differences for approximate entropy (*p* > 0.05; Fig 4). For the effect of condition, the post occlusion/pain condition exhibited increased values of the largest Lyapunov exponent and approximate entropy for both the knee and hip joint angle time series as compared to the baseline/pain free condition.

**Fig 3.**
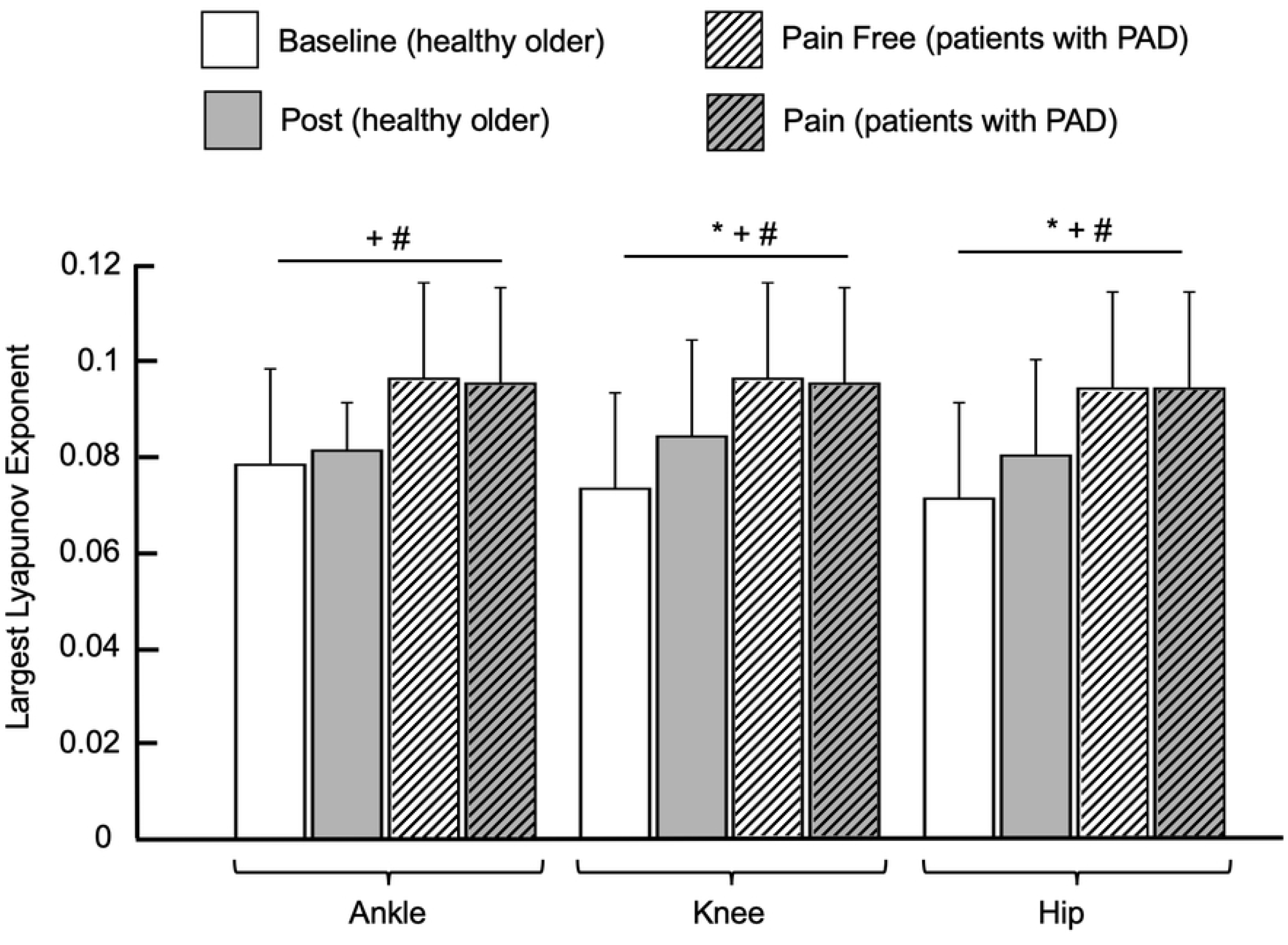
Largest Lyapunov exponent values for healthy older individuals in the baseline and post vascular occlusion (post) conditions and for patients with peripheral artery disease (PAD) in the pain free and pain conditions for ankle, knee, and hip ranges of motion. * indicates significant differences between conditions (Baseline/Pain Free vs. Post/Pain, *p* < 0.05). + indicates significant differences between groups (PAD vs. healthy older, *p* < 0.05). # indicates post-hoc tests while significant interaction exists between groups and conditions. Bar graphs represent the mean values and error bars represent the standard deviation.

**Fig 4.**
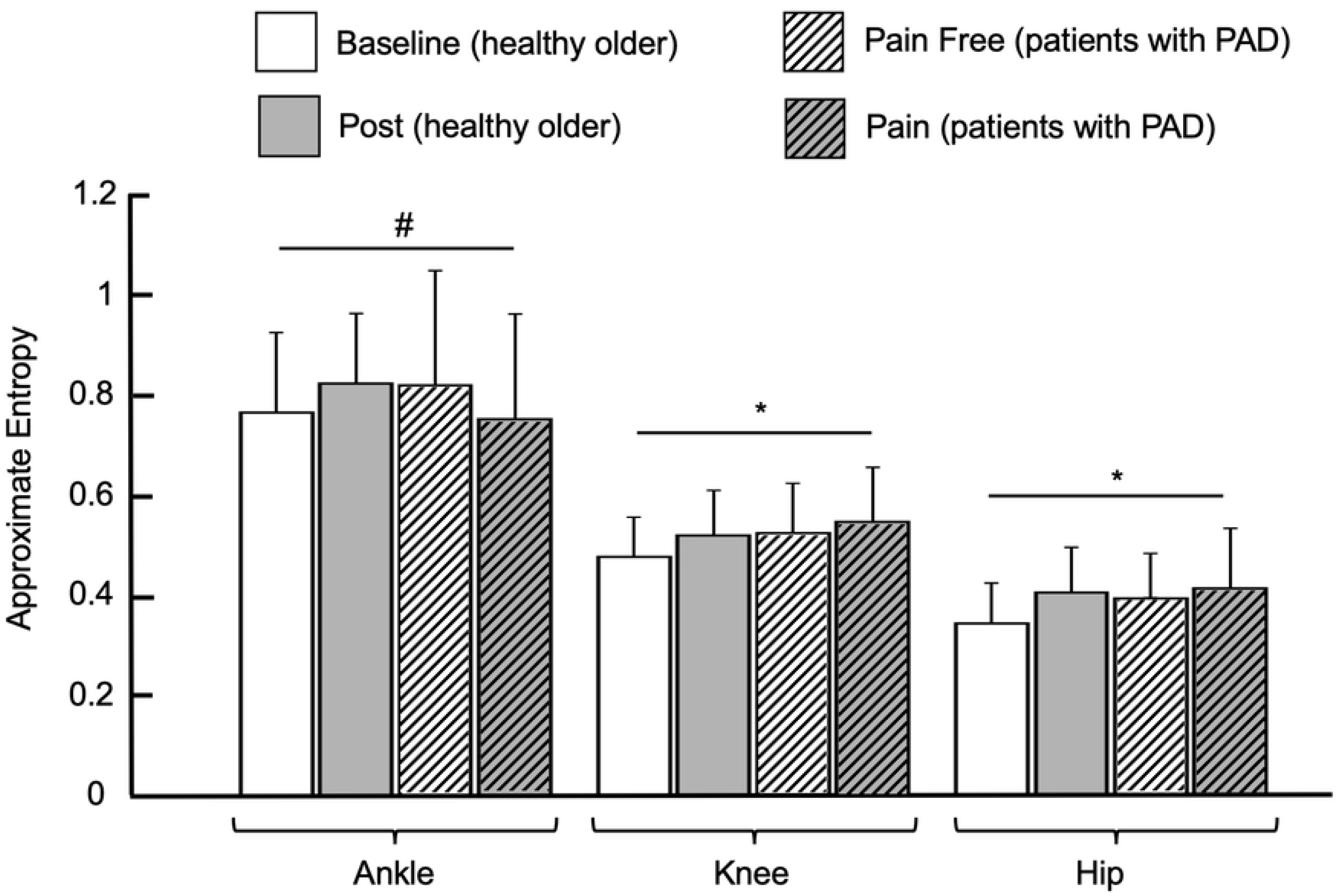
Approximate Entropy values for healthy older individuals in the baseline and post vascular occlusion (post) conditions and for patients with peripheral artery disease (PAD) in the pain free and pain conditions for ankle, knee, and hip ranges of motion. * indicates significant differences between conditions (Baseline/Pain Free vs. Post/Pain, *p* < 0.05). + indicates significant differences between groups (PAD vs. healthy older, *p* < 0.05). # indicates post-hoc tests while significant interaction exists between groups and conditions. Bar graphs represent the mean values and error bars represent the standard deviation.

There were significant interactions between group and condition for the ankle, knee, and the hip joint time series. At the ankle, the PAD pain free and PAD pain conditions demonstrated significantly increased values for the largest Lyapunov exponent and the approximate entropy of the joint angle time series as compared with the healthy control baseline and healthy control post occlusion values (Figs 3 and 4). Additionally, the approximate entropy of the ankle joint time series during post occlusion condition of the healthy controls was significantly increased compared to the baseline condition. At the knee, the largest Lyapunov exponent values of the joint angle time series were also increased in the PAD pain free and the PAD pain conditions compared to both the healthy control baseline and healthy control post occlusion conditions. The post occlusion condition in the healthy control group was also significantly increased compared to the baseline condition for the largest Lyapunov exponent values of the knee joint time series. The significant interactions at the hip were identical to those at the knee, with the PAD pain and PAD pain free conditions having significantly increased largest Lyapunov exponent values for the hip joint time series compared to the healthy control baseline and post occlusion conditions, and the healthy control post occlusion being increased compared to the baseline condition.

Results of the nonlinear analysis demonstrate that overall, patients with PAD have increased values of nonlinear analysis measures as compared to healthy older controls, regardless of the blood flow status. However, reduced blood flow also led to increases in values of measures of temporal structure of variability. The post hoc analyses further support these results as they generally show that patients with PAD have higher values of nonlinear measures during both conditions compared with both conditions in healthy older controls. These increased values indicate irregularity of the joint angle time series and increased divergence in consecutive strides of the joint movement patterns.

### 3.3 Receiver operating characteristics curve

Of the twelve individual variables, six gait variability parameters had a significant relationship with disease status and also had area under the curve values ≥ 0.75. Those variables were standard deviation of the hip joint range of motion, coefficient of variation of the knee and the hip joint ranges of motion, and the largest Lyapunov exponent at the ankle, knee, and the hip joint angle time series (Fig 5). For these six variables, area under the curve values ranged from 0.776 to 0.841, which is interpreted as “acceptable discrimination” to “excellent discrimination” of the accuracy statistic. The percent of subjects correctly identified by individual variables ranged from 71.7% to 80.0%, while sensitivity ranged from 46.7 to 90.0% and specificity ranged from 63.3 to 96.7%. Optimal cut-off points and prediction equations are also reported in Fig 5. To predict the disease status of an observation, the prediction equation is used to determine the *logit*. Then the *logit* value is used in Equation 1 to determine the probability. If the probability is greater than the cut-off point, then the observation is classified as having PAD. If the probability is less than the cut-off point, then the observation is classified as healthy. These results demonstrate that several gait variability variables provide acceptable to excellent discrimination between groups.

**Fig 5.**
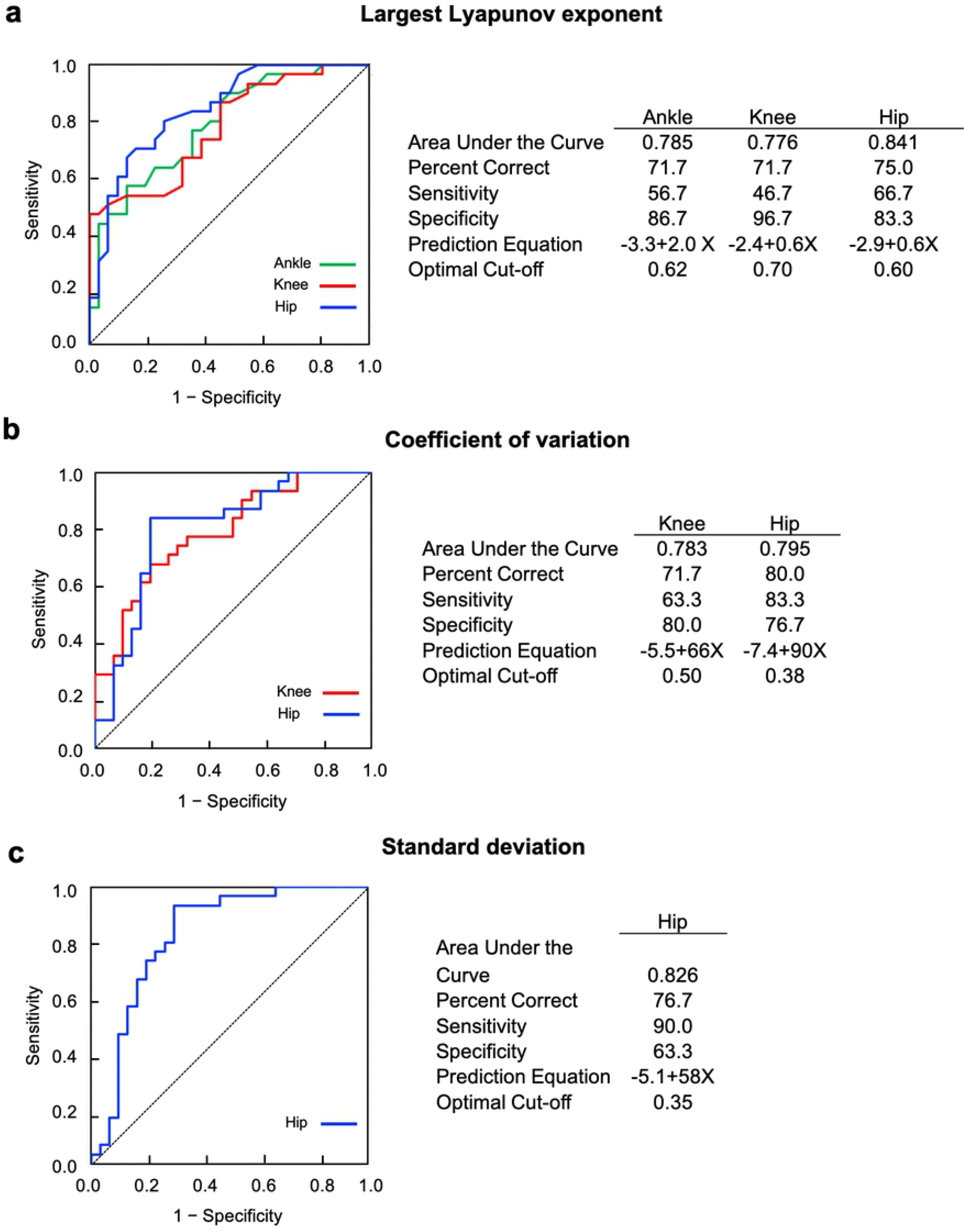
Results from receiver operating characteristics (ROC) curve and logistic regression analyses. Six dependent variables have significant relationship exist with disease status and all of them have area under the curve ≥ 0.75. a) ROC curves for Largest Lyapunov exponent of ankle, knee, and hip ranges of motion, b) ROC curves for coefficient of variation of knee and hip ranges of motion, c) ROC curve for standard deviation of hip range of motion.

## 4. Discussion

This study sought to examine the effect of reduced blood flow as the central mechanism contributing to the walking limitations seen in PAD by evaluating lower extremity gait variability in healthy older controls and patients with PAD during rested and reduced blood flow conditions. Our first hypothesis was that gait variability would be different between healthy older controls and patients with PAD. Results of our variability analyses supported this hypothesis, as the amount of variability values were significantly increased at the knee and hip ranges of motion in patients with PAD. Temporal structure of variability values were also increased at all three joint angle time series for most measures when compared with healthy older controls. Our second hypothesis was that the reduced blood flow conditions would demonstrate increased gait variability values compared with gait variability in the rested condition. Regarding this hypothesis, our results were supported at the knee and hip joints for measures of amount and temporal structure of variability. However, the ankle data did not support this, as there were no significant differences for conditions for measures of amount nor temporal structure of variability. Interestingly, significant interactions between group and condition occurred at the ankle for largest Lyapunov exponent, approximate entropy, and coefficient of variation suggesting that the ankle is affected by more than just reduced blood flow. We also analyzed our data to see if the gait variability would be similar between the healthy older control post vascular occlusion condition and the patients with PAD in the pain condition. We found that the two groups in the restricted blood flow condition had similar amount of variability but differed in the temporal structure of variability. The lack of significant interactions between the PAD pain and healthy older post occlusion conditions for the linear variability measures shows that the amount of variability is consistent between the conditions. However, the PAD pain condition had significantly increased values of the largest Lyapunov exponent for the ankle, knee, and hip joint angle time series and significantly decreased approximate entropy of the ankle joint time series as compared to the healthy older control post occlusion condition. Thus, while the variability in the amount individuals move does not seem to change, the subtle interactions in the movement patterns remain distinct between the PAD pain condition and the healthy older post occlusion condition.

An additional objective of this study was to determine the sensitivity and specificity of gait variability parameters in predicting disease status (presence or absence of PAD) using a ROC curve analysis. We hypothesized that gait variability would be able to provide good discrimination between individuals with PAD and those without PAD. The ROC curve analysis of gait variability parameters supported this hypothesis. Specifically, a combination of six gait variability parameters could correctly identify disease status of subjects 70% of the time or more. These variables included the largest Lyapunov exponent at the ankle, knee, and the hip joint time series, the standard deviation of the hip range of motion, and the coefficient of variation of the knee and the hip joint ranges of motion. Area under the curve values for these parameters indicated acceptable to excellent discrimination between groups. The majority of PAD patients are asymptomatic especially at the early stages of the disease [35], therefore, a combination of gait variability parameters could be used as a diagnostic tool for identifying patients in the community that demonstrate gait characteristics that are abnormal and suggest the development of PAD.

Gait variability has previously been utilized to study the functional limitations of patients with PAD [7,8]. Multiple components of the locomotor system must cooperate to produce gait. Variability in gait provides insight into the ability of patients with PAD to flexibly adapt to continuously changing movement environmental conditions. Gait variability is commonly used as an indicator of overall health of the biological system in older and pathological populations [54,68,69]. Previous studies found that patients with PAD have increased amount and altered temporal structure of gait variability compared with healthy controls [7,8]. Gait alterations in patients with PAD are consistently present prior to the onset of claudication pain and even from the initial steps of the patient when the oxygen levels in the legs have not been challenged yet [6–8,42]. Our current study is the first to attempt to dissect the potential contribution of reduced blood flow which many investigators feel is a key mechanism operating to produce the gait alterations of PAD. Results of the comparisons between PAD and healthy older control groups are in agreement with previous studies from our laboratory, with patients with PAD demonstrating increased amount and altered temporal structure of variability as compared with healthy controls [7,8]. The actual values of gait variability parameters are also similar as previous findings.

Previous work in healthy young also demonstrated similar post vascular occlusion values as the healthy older individuals from the current study [44]. Therefore, the current and previous studies support the notion that reduced blood flow, as induced through a vascular occlusion, contributes to alterations in the amount and structure of gait variability patterns in patients with PAD. Additionally, age does not appear to have a significant impact on the response to reduced blood flow because healthy young and older demonstrated similar responses to the vascular occlusion. Our study demonstrates that gait variability is affected by PAD and by the presence of reduced blood flow.

The existence of group differences across conditions, as indicated by several significant interactions, suggests that factors other than reduced blood flow contribute to gait variability differences seen in patients with PAD. Gait variability in the post vascular occlusion condition of healthy controls, which mimics the arterial blood flow supply-demand imbalance seen in patients with PAD, was significantly different than that of the PAD pain condition. Our first study investigating the effect of induced vascular occlusion on gait variability in healthy young forecast such a finding. That study compared the percentage change from baseline to post vascular occlusion seen in healthy young with the percentage difference from young baseline gait variability values and gait variability in patients with PAD [44]. While the post vascular occlusion condition was significantly different than the baseline condition in healthy young, the patients with PAD had significantly larger percentage differences in gait variability as compared to the increases seen in the young. For example, the percentage increases averaged across the ankle, knee, and hip for the healthy young were 21% for the largest Lyapunov exponent, 26% for the standard deviation, and 22% for the coefficient of variation. The same percent differences from healthy young baseline to the PAD pain free condition were 48% for the largest Lyapunov exponent, 62% for the standard deviation, and 99% for the coefficient of variation. This result is further supported in a similar comparison of gait variability changes from baseline to post vascular occlusion in healthy older adults. The significantly larger percentage differences in gait variability as compared to the increases seen in the young post occlusion suggest that reduced blood flow is only one mechanism contributing to differences between patients with PAD and healthy individuals. The results of the direct statistical comparison between patients with PAD and healthy older controls during rested and reduced blood flow conditions in the current study confirms and further advance the observations of the previous study. Accordingly, gait variability alterations seen in patients with PAD must be further influenced by other disease-related mechanisms, in addition to being impacted by reduced blood flow.

Maintaining an optimal “state” of gait variability is desirable because such a state represents a healthy neuromusculoskeletal system that adapts to perturbations that occur during typical movement situations. Alterations in gait variability happen because of changes in the locomotor system. This includes changes to the movement generation and control pathways of the muscular and nervous systems and the circulatory system that supplies them. There is extensive evidence of damage in the lower extremity muscles and nerves of patients with PAD, so it is likely that gait variability is also affected by these changes [34,38,39,70]. In the muscular system of patients with PAD, skeletal muscles exhibit atrophy, denervation, and defective mitochondrial bioenergetics [38–41,43,70–73]. Changes to the lower extremity nervous system in patients with PAD include axonal nerve loss as indicated by electrodiagnostic and muscle strength and control abnormalities [16,74]. How the muscular and nervous systems of patients with PAD change as the disease progresses or following treatment to restore blood flow are the obvious next steps of investigation.

An additional observation of importance includes the findings at the ankle joint time series. The current study demonstrated only one significant main effect for the ankle out of eight possible. Specifically, the PAD group has significantly increased values of the largest Lyapunov exponent as compared with controls. Several gait studies reported that the ankle joint is significantly altered in patients with PAD when compared to healthy controls [11,12,45,75]. In the current study, alterations at the ankle are apparent through multiple significant interactions. Results of the post hoc analysis revealed decreased values for the coefficient of variation of ankle range of motion and the approximate entropy of the ankle joint time series for the PAD pain condition as compared with the PAD pain free and healthy older control post vascular occlusion conditions. Therefore, the healthy older group was able to maintain the level of variability at the ankle during the post vascular occlusion condition, while patients with PAD could not. Investigation of other measures of both amount and temporal structure of variability reveals that in patients with PAD, all ankle variables decreased from the pain free to the pain condition, although not significantly. These trends of decreased values for measures of amount and temporal structure of variability of the ankle joint time series are in agreement with our previous study investigating the effect of claudication pain [45]. In that study, we suggested that patients with PAD may attempt to decrease the use of the calf muscles, the typical location for experiencing claudication pain. It is also possible that patients with PAD are so limited during claudication pain that even the abnormal movement patients exhibited during pain free walking is not possible to maintain. Future studies could investigate these hypotheses by assessing changes in calf muscle activity before and after the onset of claudication pain.

Results of the receiver operating characteristics curve analysis demonstrate that multiple gait variability parameters have potential for predicting PAD disease status (presence or absence of PAD). Interestingly, three of the four dependent variables assessing variability of hip joint movement were significantly related with disease status. The hip variability measures also predicted disease status most correctly among the six significant gait variability parameters. The largest Lyapunov exponent for the ankle, knee, and the hip joint time series were significantly related with disease status. This suggests that this nonlinear measure is more strongly associated with disease status, regardless of the joint being observed.

Receiver operating characteristics curve analysis has been used extensively to predict lower extremity injuries, including falling in the older adults [64,76–78]. In comparison with previous studies using functional tests to identify individuals at risk for falls, the current study has equal or better values of sensitivity and specificity. For example, Thrane et al. examined the predictive value of the timed up and go test for determining fall risk [76]. Sensitivity values in this study ranged from 11 to 44% and specificity varied from 58 to 98%. Sensitivity in our study went from 47 to 90% and specificity ranged from 63 to 97%. Therefore, our results demonstrate that gait variability is a sensitive and specific assessment of the lower extremity mobility problems experienced by patients with PAD. Although several variables demonstrated a significant relationship with disease status, future studies should explore multivariate analysis and machine learning models, as the combination of multiple gait variability parameters might yield even better diagnostic capabilities.

The difficulty in interpreting the results of our study lies in the classifications of our subject conditions. Our study sought to determine the effect of reduced blood flow on gait variability alterations that occur in patients with PAD. Therefore, we examined both patients with PAD and healthy older controls during rested and reduced blood flow conditions. Both groups had lower extremity blood flow assessed at rest and we acknowledge that patients with PAD have abnormal blood flow even at rest. Similarly, we acknowledge that the vascular occlusion procedure performed to reduce lower extremity blood flow in the healthy older group did not lead to hemodynamics identical to those experienced by patients with PAD. However, the approach was effective in producing perturbed blood flow and significant alterations in gait during the post vascular occlusion condition. Furthermore, the reductions in blood flow delivery from a rested state to an exerted state, documented by the ankle-brachial indices during rest and exercise in patients with PAD [79,80], are reflective of the lower extremity hemodynamics during the PAD pain free (rested) and PAD pain (exercise) conditions.

Although it was not possible to continuously monitor the ankle-brachial index of healthy older controls during the post vascular occlusion condition, the current protocol created a critical episode of clinical ischemia consistent with reduced arterial inflow. The vascular occlusion procedure was modeled after the vascular laboratory technique for inducing ischemia and reperfusion, which creates a supply-demand imbalance similar to that experienced by patients with PAD [46–51]. Therefore, our study provides important insights into the effect of reduced blood flow on gait variability parameters, regardless of the difficulties in simulating the blood flow status of PAD in healthy individuals. The vascular occlusion method used in our study could be validated by newly available technologies, such as a tissue oximeter, to better document blood flow delivery to the calf muscles during rested and post vascular occlusion conditions.

## 5. Conclusions

The results of our study overall demonstrate that gait variability is affected both by PAD and by the induction of an episode of reduced blood flow. The presence of group differences across conditions supports the findings of our previous studies that a number of other factors, in addition to reduced blood flow, contribute to gait variability differences between healthy individuals and those with PAD. Other potential mechanisms include the documented damaged muscle structure, changes in muscle bioenergetics, and nerve loss in patients with PAD. These gait variability changes in healthy older individuals caused by the induced vascular occlusion should highlight the importance of screening older individuals at risk of developing PAD, as potential mobility problems and altered gait variability can develop with reductions of blood flow, even if no claudication symptoms exist. Gait variability parameters have potential diagnostic value, as some gait variability measures had 90.0% probability of identifying patients with PAD. Future research should investigate this further, including incorporating multivariate prediction models to more accurately classify patients with PAD and healthy individuals.

## References

1. Nicoloff AD, Taylor J, McLafferty RB, Moneta GL, Porter JM, Menzoian JO. Patient recovery after infrainguinal bypass grafting for limb salvage. J Vasc Surg. 1998;27: 256–266. doi:10.1016/S0741-5214(98)70356-8

2. Toursarkissian B, Shireman PK, Harrison A, D’Ayala M, Schoolfield J, Sykes MT. Major lower-extremity amputation: Contemporary experience in a single Veterans Affairs institution. Am Surg. 2002;68: 606–610.

3. Jaffery Z, Greenbaum AB, Siddiqui MF, Mahendraker N, Gupta V, Mokkala V, et al. Predictors of mortality in patients with lower extremity peripheral arterial disease: 5-year follow-up. J Interv Cardiol. 2009;22: 564–570. doi:10.1111/j.1540-8183.2009.00505.x

4. Gardner AW, Montgomery PS. Impaired balance and higher prevalence of falls in subjects with intermittent claudication. J Gerontol A Biol Sci Med Sci. 2001;56: M454–8. doi:10.1093/gerona/56.7.m454

5. Gardner AW, Montgomery PS. The relationship between history of falling and physical function in subjects with peripheral arterial disease. Vasc Med. 2001;6: 223–227. doi:10.1177/1358836x0100600404

6. Hernandez H, Myers SA, Schieber M, Ha DM, Baker S, Koutakis P, et al. Quantification of Daily Physical Activity and Sedentary Behavior of Claudicating Patients. Ann Vasc Surg. 2019;55: 112–121. doi:10.1016/j.avsg.2018.06.017

7. Myers SA, Pipinos II, Johanning JM, Stergiou N. Gait variability of patients with intermittent claudication is similar before and after the onset of claudication pain. Clin Biomech. 2011;26: 729–734. doi:10.1016/j.clinbiomech.2011.03.005

8. Myers SA, Johanning JM, Stergiou N, Celis RI, Robinson L, Pipinos II. Gait variability is altered in patients with peripheral arterial disease. J Vasc Surg. 2009;49: 924–931.e1. doi:10.1016/j.jvs.2008.11.020

9. Chen SJ, Pipinos I, Johanning J, Radovic M, Huisinga JM, Myers SA, et al. Bilateral claudication results in alterations in the gait biomechanics at the hip and ankle joints. J Biomech. 2008;41: 2506–2514. doi:10.1016/j.jbiomech.2008.05.011

10. Celis R, Pipinos II, Scott-Pandorf MM, Myers SA, Stergiou N, Johanning JM. Peripheral arterial disease affects kinematics during walking. J Vasc Surg. 2009;49: 127–132. doi:10.1016/j.jvs.2008.08.013

11. Koutakis P, Johanning JM, Haynatzki GR, Myers SA, Stergiou N, Longo GM, et al. Abnormal joint powers before and after the onset of claudication symptoms. J Vasc Surg. 2010;52: 340–347. doi:10.1016/j.jvs.2010.03.005

12. Koutakis P, Pipinos II, Myers SA, Stergiou N, Lynch TG, Johanning JM. Joint torques and powers are reduced during ambulation for both limbs in patients with unilateral claudication. J Vasc Surg. 2010;51: 80–88. doi:10.1016/j.jvs.2009.07.117

13. Makris KI, Nella AA, Zhu Z, Swanson SA, Casale GP, Gutti TL, et al. Mitochondriopathy of peripheral arterial disease. Vascular. 2007;15: 336–343. doi:10.2310/6670.2007.00054

14. Cluff K, Miserlis D, Naganathan GK, Pipinos II, Koutakis P, Samal A, et al. Morphometric analysis of gastrocnemius muscle biopsies from patients with peripheral arterial disease: Objective grading of muscle degeneration. Am J Physiol - Regul Integr Comp Physiol. 2013;305. doi:10.1152/ajpregu.00525.2012

15. Shu J, Santulli G. Update on peripheral artery disease: Epidemiology and evidence-based facts. Atherosclerosis. Elsevier Ireland Ltd; 2018. pp. 379–381. doi:10.1016/j.atherosclerosis.2018.05.033

16. Schieber MN, Hasenkamp RM, Pipinos II, Johanning JM, Stergiou N, DeSpiegelaere HK, et al. Muscle strength and control characteristics are altered by peripheral artery disease. Journal of Vascular Surgery. Mosby Inc.; 2017. pp. 178–186.e12. doi:10.1016/j.jvs.2017.01.051

17. Tinetti ME, Doucette JT, Claus EB. The Contribution of Predisposing and Situational Risk Factors to Serious Fall Injuries. J Am Geriatr Soc. 1995;43: 1207–1213. doi:10.1111/j.1532-5415.1995.tb07395.x

18. Callisaya ML, Blizzard L, Schmidt MD, Martin KL, McGinley JL, Sanders LM, et al. Gait, gait variability and the risk of multiple incident falls in older people: a population-based study. Age Ageing. 2011;40: 481–487. doi:10.1093/ageing/afr055

19. Hausdorff JM, Rios DA, Edelberg HK. Gait variability and fall risk in community-living older adults: A 1-year prospective study. Arch Phys Med Rehabil. 2001;82: 1050–1056. doi:10.1053/apmr.2001.24893

20. Glass L, Mackey MC. From clocks to chaos: the rhythms of life. Princeton University Press; 1988.

21. Antonellis P, Galle S, De Clercq D, Malcolm P. Altering gait variability with an ankle exoskeleton. PLoS One. 2018;13. doi:10.1371/journal.pone.0205088

22. Goldberger AL, Amaral LAN, Hausdorff JM, Ivanov PC, Peng CK, Stanley HE. Fractal dynamics in physiology: Alterations with disease and aging. Proc Natl Acad Sci U S A. 2002;99: 2466–2472. doi:10.1073/pnas.012579499

23. Lipsitz LA, Goldberger AL. Loss of ‘Complexity’ and Aging: Potential Applications of Fractals and Chaos Theory to Senescence. JAMA J Am Med Assoc. 1992;267: 1806–1809. doi:10.1001/jama.1992.03480130122036

24. Lipsitz LA. Dynamics of stability: The physiologic basis of functional health and frailty. Journals Gerontol - Ser A Biol Sci Med Sci. 2002;57: B115–25. doi:10.1093/gerona/57.3.B115

25. Stergiou N, Moraiti C, Giakas G, Ristanis S, Georgoulis AD. The effect of the walking speed on the stability of the anterior cruciate ligament deficient knee. Clin Biomech. 2004;19: 957–963. doi:10.1016/j.clinbiomech.2004.06.008

26. Georgoulis AD, Moraiti C, Ristanis S, Stergiou N. A novel approach to measure variability in the anterior cruciate ligament deficient knee during walking: The use of the approximate entropy in orthopaedics. J Clin Monit Comput. 2006;20: 11–18. doi:10.1007/s10877-006-1032-7

27. Hausdorff JM, Cudkowicz ME, Firtion R, Wei JY, Goldberger AL. Gait variability and basal ganglia disorders: stride-to-stride variations of gait cycle timing in Parkinson’s disease and Huntington’s disease [published erratum appears in Mov Disord 1998 Jul;13(4):757]. Mov Disord. 1998;13: 428–437.

28. Richardson JK, Thies SB, DeMott TK, Ashton-Miller JA. Gait analysis in a challenging environment differentiates between fallers and nonfallers among older patients with peripheral neuropathy. Arch Phys Med Rehabil. 2005;86: 1539–1544. doi:10.1016/j.apmr.2004.12.032

29. Webster KE, Merory JR, Wittwer JE. Gait variability in community dwelling adults with Alzheimer disease. Alzheimer Dis Assoc Disord. 2006;20: 37–40. doi:10.1097/01.wad.0000201849.75578.de

30. Ellis RJ, Ng YS, Zhu S, Tan DM, Anderson B, Schlaug G, et al. A validated smartphone-based assessment of gait and gait variability in Parkinson’s disease. PLoS One. 2015;10. doi:10.1371/journal.pone.0141694

31. Gilat M, Shine JM, Bolitho SJ, Matar E, Kamsma YPT, Naismith SL, et al. Variability of Stepping during a Virtual Reality Paradigm in Parkinson’s Disease Patients with and without Freezing of Gait. PLoS One. 2013;8. doi:10.1371/journal.pone.0066718

32. Bakir MS, Gruschke F, Taylor WR, Haberl EJ, Sharankou I, Perka C, et al. Temporal but Not Spatial Variability during Gait Is Reduced after Selective Dorsal Rhizotomy in Children with Cerebral Palsy. PLoS One. 2013;8. doi:10.1371/journal.pone.0069500

33. Rose GA. The diagnosis of ischaemic heart pain and intermittent claudication in field surveys. Bull World Health Organ. 1962;27: 645–658. Available: http://www.ncbi.nlm.nih.gov/pubmed/13974778

34. Brass EP, Hiatt WR. Acquired skeletal muscle metabolic myopathy in atherosclerotic peripheral arterial disease. Vascular Medicine. Arnold; 2000. pp. 55–59. doi:10.1177/1358836X0000500109

35. McDermott MMG, Greenland P, Liu K, Guralnik JM, Criqui MH, Dolan NC, et al. Leg symptoms in peripheral arterial disease associated clinical characteristics and functional impairment. J Am Med Assoc. 2001;286: 1599–1606. doi:10.1001/jama.286.13.1599

36. Pipinos II, Boska MD, Shepard AD, Anagnostopoulos P V., Katsamouris A. Pentoxifylline reverses oxidative mitochondrial defect in claudicating skeletal muscle. J Surg Res. 2002;102: 126–132. doi:10.1006/jsre.2001.6292

37. Pipinos II, Judge AR, Zhu Z, Selsby JT, Swanson SA, Johanning JM, et al. Mitochondrial defects and oxidative damage in patients with peripheral arterial disease. Free Radic Biol Med. 2006;41: 262–269. doi:10.1016/j.freeradbiomed.2006.04.003

38. Pipinos II, Judge AR, Selsby JT, Zhu Z, Swanson SA, Nella AA, et al. The myopathy of peripheral arterial occlusive disease: Part 1. Functional and histomorphological changes and evidence for mitochondrial dysfunction. Vascular and Endovascular Surgery. 2008. pp. 481–489. doi:10.1177/1538574407311106

39. Pipinos II, Judge AR, Selsby JT, Zhu Z, Swanson SA, Nella AA, et al. The myopathy of peripheral arterial occlusive disease: Part 2. Oxidative stress, neuropathy, and shift in muscle fiber type. Vascular and Endovascular Surgery. 2008. pp. 101–112. doi:10.1177/1538574408315995

40. Koutakis P, Miserlis D, Myers SA, Kim JKS, Zhu Z, Papoutsi E, et al. Abnormal Accumulation of Desmin in Gastrocnemius Myofibers of Patients with Peripheral Artery Disease: Associations with Altered Myofiber Morphology and Density, Mitochondrial Dysfunction and Impaired Limb Function. J Histochem Cytochem. 2015;63: 256–269. doi:10.1369/0022155415569348

41. Koutakis P, Myers SA, Cluff K, Ha DM, Haynatzki G, McComb RD, et al. Abnormal myofiber morphology and limb dysfunction in claudication. J Surg Res. 2015;196: 172–179. doi:10.1016/j.jss.2015.02.011

42. Fuglestad MA, Hernandez H, Gao Y, Ybay H, Schieber MN, Brunette KE, et al. A low-cost, wireless near-infrared spectroscopy device detects the presence of lower extremity atherosclerosis as measured by computed tomographic angiography and characterizes walking impairment in peripheral artery disease. Journal of Vascular Surgery. Mosby Inc.; 2020. pp. 946–957. doi:10.1016/j.jvs.2019.04.493

43. Ha DM, Carpenter LC, Koutakis P, Swanson SA, Zhu Z, Hanna M, et al. Transforming growth factor-beta 1 produced by vascular smooth muscle cells predicts fibrosis in the gastrocnemius of patients with peripheral artery disease. J Transl Med. 2016;14. doi:10.1186/s12967-016-0790-3

44. Myers SA, Stergiou N, Pipinos II, Johanning JM. Gait variability patterns are altered in healthy young individuals during the acute reperfusion phase of ischemia-reperfusion. J Surg Res. 2010;164: 6–12. doi:10.1016/j.jss.2010.04.030

45. Myers SA, Johanning JM, Pipinos II, Schmid KK, Stergiou N. Vascular occlusion affects gait variability patterns of healthy younger and older individuals. Ann Biomed Eng. 2013;41: 1692–1702. doi:10.1007/s10439-012-0667-4

46. Diener HC, Dichgans J, Guschlbauer B, Mau H. The significance of proprioception on postural stabilization as assessed by ischemia. Brain Res. 1984;296: 103–109. doi:10.1016/0006-8993(84)90515-8

47. Haouzi P, Huszczuk A, Porszasz J, Chalon B, Wasserman K, Whipp BJ. Femoral vascular occlusion and ventilation during recovery from heavy exercise. Respir Physiol. 1993;94: 137–150. doi:10.1016/0034-5687(93)90043-a

48. Kjaer M, Pott F, Mohr T, Linkis P, Tornøe P, Secher NH. Heart rate during exercise with leg vascular occlusion in spinal cord-injured humans. J Appl Physiol. 1999;86: 806–811. doi:10.1152/jappl.1999.86.3.806

49. Sargeant AJ, Rouleau MY, Sutton JR, Jones NL. Ventilation in exercise studied with circulatory occlusion. J Appl Physiol Respir Environ Exerc Physiol. 1981;50: 718–723. doi:10.1152/jappl.1981.50.4.718

50. Stanley WC, Lee WR, Brooks GA. Ventilation studied with circulatory occlusion during two intensities of exercise. Eur J Appl Physiol Occup Physiol. 1985;54: 269–77. doi:10.1007/bf00426144

51. Tokizawa K, Mizuno M, Nakamura Y, Muraoka I. Venous occlusion to the lower limb attenuates vasoconstriction in the nonexercised limb during posthandgrip muscle ischemia. J Appl Physiol. 2004;96: 981–984. doi:10.1152/japplphysiol.00695.2003

52. Barrett R, Noordegraaf MV, Morrison S. Gender differences in the variability of lower extremity kinematics during treadmill locomotion. J Mot Behav. 2008;40: 62–70. doi:10.3200/JMBR.40.1.62-70

53. Keenan S, Stergiou N. The reliability of the Lyapunov exponent during treadmill walking. Proceedings of the Fourth World Congress of Biomechanics Meeting. Calgary, Alberta, Canada; 2002.

54. Stergiou N, Buzzi UH, Kurz MJ, Heidel J. Nonlinear tools in human movement. In: Stergiou N, editor. Innovative analysis of human movement. Champaign, IL: Human Kinetics; 2004. pp. 63–90.

55. Abarbanel HDI. Analysis of observed chaotic data. New York: Springer-Verlag; 1996.

56. Buzzi UH, Stergiou N, Kurz MJ, Hageman PA, Heidel J. Nonlinear dynamics indicates aging affects variability during gait. Clin Biomech. 2003;18: 435–443. doi:10.1016/S0268-0033(03)00029-9

57. Wolf A, Swift JB, Swinney HL, Vastano JA. Determining Lyapunov exponents from a time series. Phys D Nonlinear Phenom. 1985;16: 285–317. doi:10.1016/0167-2789(85)90011-9

58. Sprott J, Rowlands G. Chaos data analyzer: The professional version. New York: American Institute of Physics; 1992.

59. Pincus SM, Goldberger AL. Physiological time-series analysis: What does regularity quantify? Am J Physiol. 1994;266 (4 Pt. doi:10.1152/ajpheart.1994.266.4.h1643

60. Pincus SM, Gladstone IM, Ehrenkranz RA. A regularity statistic for medical data analysis. J Clin Monit. 1991;7: 335–345. doi:10.1007/BF01619355

61. Pincus SM. Approximate entropy (ApEn) as a regularity measure. In: Newell KM, Molenaar PCM, editors. Applications of nonlinear dynamics to developmental process modeling. Mahwah, NJ: Lawrence Erlbaum Associates; 1998. pp. 243–268.

62. Pincus S. Approximate entropy (ApEn) as a complexity measure. Chaos. 1995;5: 110–117. doi:10.1063/1.166092

63. Pincus SM. Irregularity and asynchrony in biologic network signals. Methods in enzymology. 2000. pp. 149–182. doi:10.1016/s0076-6879(00)21192-0

64. Bruce S, Wilkerson G. Clinical prediction rules, part 1: Conceptual overview. Athl Ther Today. 2010;15: 4–9.

65. Fawcett T. An introduction to ROC analysis. Pattern Recognit Lett. 2006;27: 861–874. doi:10.1016/j.patrec.2005.10.010

66. Hosmer DW, Lemeshow S. Assessing the Fit of the Model. In: Applied Logistic Regression. John Wiley and Sons, Inc. John Wiley & Sons, Inc.; 2000. doi:10.1002/0471722146

67. Zar J. Biostatistical Analysis. Fifth. Upper Saddle River, New Jersey: Prentice Hall Inc.; 2010.

68. Hausdorff JM. Gait variability: Methods, modeling and meaning. Journal of NeuroEngineering and Rehabilitation. 2005. doi:10.1186/1743-0003-2-19

69. Hausdorff JM. Gait dynamics, fractals and falls: Finding meaning in the stride-to-stride fluctuations of human walking. Hum Mov Sci. 2007;26: 555–589. doi:10.1016/j.humov.2007.05.003

70. McDermott MM, Hoff F, Ferrucci L, Pearce WH, Guralnik JM, Tian L, et al. Lower extremity ischemia, calf skeletal muscle characteristics, and functional impairment in peripheral arterial disease. J Am Geriatr Soc. 2007;55: 400–406. doi:10.1111/j.1532-5415.2007.01092.x

71. Regensteiner JG, Wolfel EE, Brass EP, Carry MR, Ringel SP, Hargarten ME, et al. Chronic changes in skeletal muscle histology and function in peripheral arterial disease. Circulation. 1993;87: 413–421. doi:10.1161/01.CIR.87.2.413

72. Becker RA, Cluff K, Duraisamy N, Mehraein H, Farhoud H, Collins T, et al. Optical probing of gastrocnemius in patients with peripheral artery disease characterizes myopathic biochemical alterations and correlates with stage of disease. Physiol Rep. 2017;5. doi:10.14814/phy2.13161

73. Koutakis P, Weiss DJ, Miserlis D, Shostrom VK, Papoutsi E, Ha DM, et al. Oxidative damage in the gastrocnemius of patients with peripheral artery disease is myofiber type selective. Redox Biol. 2014;2: 921–928. doi:10.1016/j.redox.2014.07.002

74. Weber F, Ziegler A. Axonal neuropathy in chronic peripheral arterial occlusive disease. Muscle and Nerve. 2002;26: 471–476. doi:10.1002/mus.10235

75. Crowther RG, Spinks WL, Leicht AS, Quigley F, Golledge J. Relationship between temporal-spatial gait parameters, gait kinematics, walking performance, exercise capacity, and physical activity level in peripheral arterial disease. J Vasc Surg. 2007;45: 1172–1178. doi:10.1016/j.jvs.2007.01.060

76. Thrane G, Joakimsen RM, Thornquist E. The association between timed up and go test and history of falls: The Tromso study. BMC Geriatr. 2007;7: 1–7. doi:10.1186/1471-2318-7-1

77. Tzagarakis GN, Tsivgoulis SD, Papagelopoulos PJ, Mastrokalos DS, Papadakis NC, Kampanis NA, et al. Influence of acute anterior cruciate ligament deficiency in gait variability. J Int Med Res. 2010;38: 511–525. doi:10.1177/147323001003800214

78. Wrisley DM, Kumar NA. Functional Gait Assessment: Concurrent, Discriminative, and Predictive Validity in Community-Dwelling Older Adults. Phys Ther. 2010;90: 761–773. doi:10.2522/ptj.20090069

79. Skinner JS, Strandness DE. Exercise and intermittent claudication. I. Effect of repetition and intensity of exercise. Circulation. 1967;36: 15–22. doi:10.1161/01.cir.36.1.15

80. Stahler C, Strandness DE. Ankle Blood Pressure Response to Graded Treadmill Exercise. Angiology. 1967;18: 237–241. doi:10.1177/000331976701800406

